# Genetic evidence indicates new distribution record of endangered Kashmir musk deer (*Moschus cupreus*) with range expansion in Uttarakhand, India

**DOI:** 10.1101/2020.08.20.258962

**Authors:** Ajit Kumar, Bhim Singh, Subhashree Sahoo, Kumudani Bala Gautam, Sandeep Kumar Gupta

## Abstract

Kashmir musk deer, KMD (*Moschus cupreus*) is one the most threatened species reported from the Himalayan region of Kashmir, Pakistan and Afghanistan. A comprehensive and reliable distribution range of musk deer is still lacking. Recently, a molecular study confirmed the presence of KMD in Mustang in Nepal, west of Annapurna Himalayas. Here, we investigated the phylogenetic relationship of musk deer from Jammu and Kashmir (J&K), Kedarnath Wildlife Sanctuary (KWLS), and Nanda Devi Biosphere Reserve (NDBR), Uttarakhand region, India based on mitochondrial control region. The Bayesian phylogenetic analysis indicated a close genetic relationship between samples from J&K, KWLS and NDBR with identified lineages of KMD from Nepal with high posterior probabilities (PP∼100). It confirmed that the musk deer lineage from the Uttarakhand region of KWLS (1025-3662 m) and NDBR (1800-7817 m) to be of KMD (*M. cupreus*) and hence a distinct Evolutionary Significant Unit (ESU). Besides, as per the IUCN database, the Western Himalayan region also holds the population of *M. leucogaster* and *M. chrysogaster*. Hence, we suggest extensive sampling for proper identification and validation of the geographic limits of musk deer species. We report for the first time the existence of KMD from the Uttarakhand region that we recommend to be updated in the IUCN database. It will assist in the effective conservation and management of this enigmatic endangered species.

## Introduction

Musk deer belongs to the genus *Moschus* in the monotypic family Moschidae. The species are endemic to the Palearctic region mostly inhabiting fragmented geographical areas of Indian Himalaya, Tibetan Plateau and its adjoining mountainous region in China and the Far East (Pan et al., 2015). Musk deer are habitat specialist solitary animals found in the alpine shrubland and above the treeline of alpine meadows at an altitude of 2600-4600m. At present, seven species of musk deer are recognized, of which, five species: Kashmir musk deer (*M. cupreus*), Alpine musk deer (*M. chrysogaster*), Himalayan musk deer (*M. leucogaster*), Forest musk deer (*M. berezovskii*) and Black musk deer (*M. fuscus*) are found in the Indian Himalayan range (Grubb 2005). The populations of all musk deer are dwindling due to heavy poaching for musk pod and habitat fragmentation and degradation due to anthropogenic pressure. Due to unsustainable exploitation, all musk deer have been included in Appendices of Convention on International Trade in Endangered Species of Wild Fauna and Flora (CITES) since 1979 (Zhou et al., 2004). The International Union for Conservation of Nature (IUCN) Red data list enlists six species of musk deer under the ‘Endangered’ category while one species is in the ‘Vulnerable’ category. In India, musk deer are included in Schedule *I* under the Indian Wild Life (Protection) Act, 1972 (WPA). Due to overlapping distribution ranges, there is ambiguity in their species taxonomy impeding efficient conservation efforts (Pang et al. 2015).

The Kashmir musk deer (KMD) is one of the least studied species of musk deer. Previously, KMD has been reported from the Himalayan region of Kashmir, Pakistan and Afghanistan (Grubb 2005). Due to limited baseline information on ecological and genetic data, the actual distribution range of KMD is still not resolved. However, a recent molecular and camera-trap based study reported a new distribution record of KMD from Mustang in Nepal, west of Annapurna Himalayas range (Singh et al., 2019). Musk deer is highly cryptic, which makes species validation solely based on morphological characteristics unreliable (Su et al., 2001; Groves et al., 1994; Groves et al., 1987). Moreover, the Alpine and the Himalayan musk deer appear very similar to KMD with coat color undergoing seasonal variation (Liu, and Groves, 2014, Singh et al., 2019). The use of advanced molecular tools for species identification and phylogenetic analysis has led to the resolution of phylogenetic complexities in musk deer and aided species validation (Pan et al., 2015, Su et al., 2001). Genetic studies were a vital resource that confirmed the presence of Himalayan musk deer, which was misidentified as Alpine musk deer in Tibet (Guo *et al*., 2018) and the presence of Eastern lineages of hog deer (*Axis porcinus annamiticus*) from the Keibul Lamjao National Park, Manipur India (Gupta et al., 2018). The mitochondrial DNA (mtDNA) control region (CR) has proven to be a powerful marker for investigating the intra-species genetic variation (Hu et al. 2006, Peng et al., 2008, Kumar et al., 2017; Gupta et al., 2018). The KWLS is one of the largest Protected Areas in the Western Himalaya, Uttarakhand covering a high altitudinal area of 975 km^2^. In the eastern part of Kedarnath Wildlife Sanctuary (KWLS), the Valley of Flowers national park forms a part of the Nanda Devi Biosphere Reserve (NDBR). We aimed to examine the phylogenetics among the musk deer samples collected from the Himalayan regions of Jammu and Kashmir (J&K) and Uttarakhand (UK) to confirm the species and furnishing baseline information for the molecular forensics.

## Methodology

### Sample collection and DNA extraction

We used 20 biological samples (musk pod, tissue, and hair) of musk deer from KWLS (n=18), NDBR (n=1), from the UK and one sample of KMD from Srinagar, J&K sent by the State Forest Department (Fig. 1). The samples were preserved at -80°C until DNA extraction. We extracted genomic DNA (gDNA) from the samples using a modified DNeasy Blood & Tissue kit (Qiagen, Hilden, Germany) protocol. The authors confirm that all the experiments were performed following relevant guidelines and regulations.

**Figure 1:**
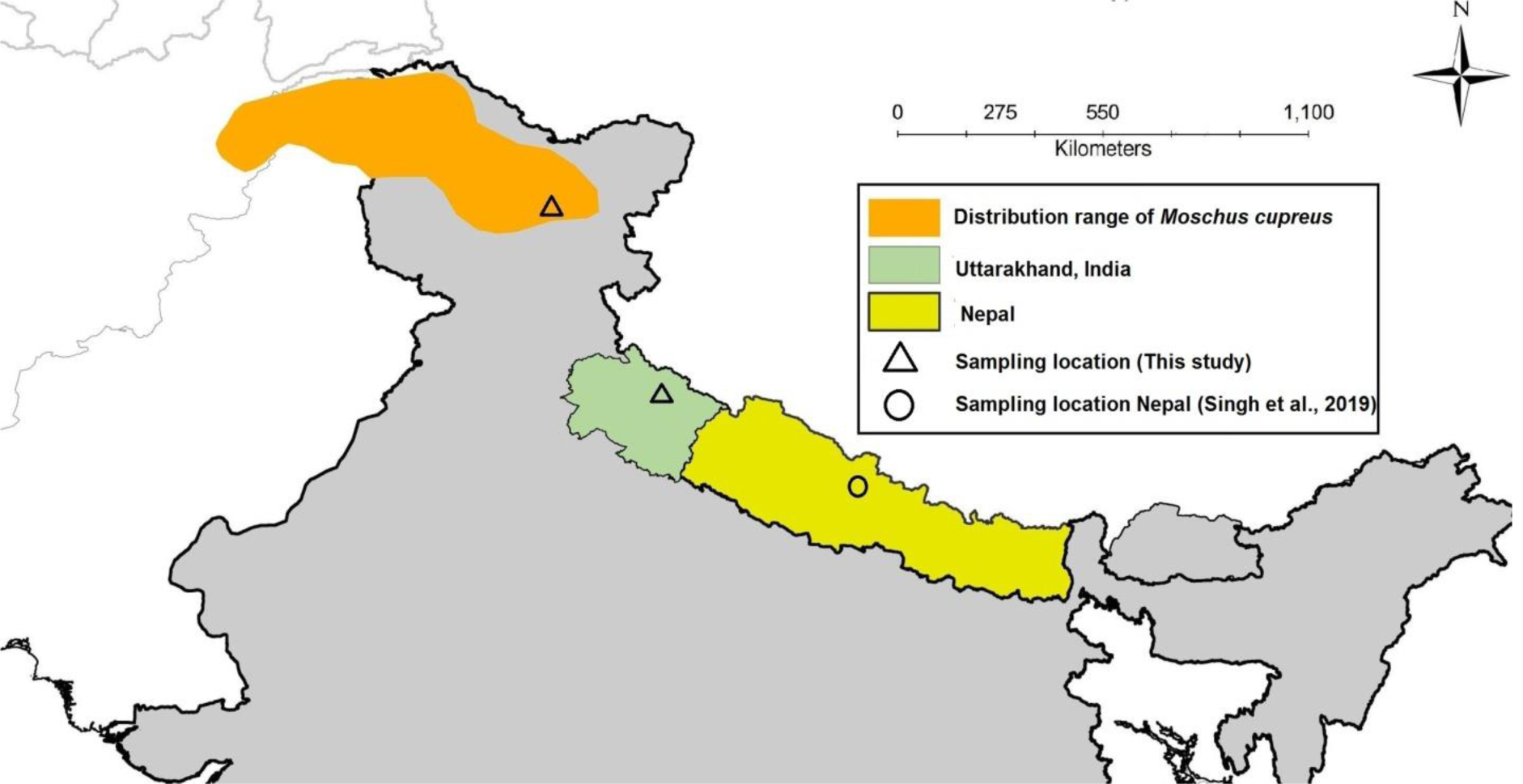
Sampling location and distribution range of Kashmir musk deer (*Moschus cupreus*) as per the IUCN record.

### PCR amplification and sequencing

The reactions were performed in 20μl volumes containing 10-20 ng of extracted genomic DNA. PCR master mix contained: 1× PCR buffer (Applied Biosystem), 2.0 mM MgCl2, 0.2 mM of each dNTP, 2 pmol of each primer, and 5U of Taq DNA polymerase. We successfully amplified 485 bp long portions of mtDNA CR using Cerv.tPro: 5’-CCACYATCAACACCCAAAGC-3’; CervCRH: 5’-GCCCTGAARAAAGAACCAGATG-3’ (Balakrishnan et al., 2003). The PCR conditions for both the primer were as follows: an initial denaturation for 5 minutes at 95°C, followed by 35 cycles at 95°C for 45 seconds, 55°C for 45 seconds and 72°C for 45 seconds, with a final extension of 72°C for 15 minutes. The efficiency and reliability of PCR reactions were monitored by using control reactions. The PCR products were electrophoresed on 2% agarose gel and visualized under UV light. Positive amplicons were treated with Exonuclease-I and Shrimp alkaine phosphatase (USB, Cleveland, OH) for 15 min each at 37°C and 80°C respectively to remove any reaction residues. The purified fragments were sequenced directly in Applied Biosystems Genetic Analyzer 3500XL from both primers set using a BigDye v3.1 Kit.

### Data Analysis

The generated sequences were obtained from both directions of targeted mtDNA fragments which then were edited with SEQUENCHER® version 4.9 (Gene Codes Corporation, Ann Arbor, MI, USA). CLUSTAL *X* 1.8 multiple alignment program was used to aligned all sequences and alignment was checked by visual inspection (Thompson et al., 1997). DnaSP 5.0 (Librado et al.,2009) was used to calculate the number of haplotypes in the data set. For the phylogenetic reconstruction, we included the sequences of *M. moschiferus* (n=2), *M. chrysogaster* (n=3), *M. anhuiensis* (n=1), *M. leucogaster* (n=20), *M. berezovskii* (n=24) and *M. cupreus* (n=5) from GenBank (Supplementary Table: ST1). Bayesian Inference (BI) of the phylogenetic relationship among all sequences of mtDNA CR was constructed by using BEAUti v 1.6.1 and BEAST v.1.10.4 (Drummond et al. 2012). One sequence of Indian Mouse deer (*Moschiola indica*) (NC037993) was chosen as the outgroup. We deployed best-fit nucleotide substitution model Hasegawa–Kishono–Yano (HKY)+G+I to obtain the best tree topology in phylogenetic analysis. Bayesian inference analysis was run for four simultaneous MCMC chains for 10 million generations and sampled every 100 generations using a burn-in of 5000 generations. The resulting phylogenetic trees were visualized in FigTree v1.4.0 (http://tree.bio.ed.ac.uk/software/figtree/).

The evolutionary divergence over sequence pairs between musk deer groups was estimated with a p-distance model including both substitution transition and transversion calculated in MEGA X (Kumar et al., 2018).

## Results and Discussion

The generated 20 sequences of mtDNA CR region of musk deer species from the present study were compared with previously published sequenced (n=5) of KMD from Nepal. All 25 sequences were grouped into 8 haplotypes (Hap). Out of the 8 haplotypes, Hap 1 was common in both Nepal and Uttarakhand populations representing three and seven sequences, respectively. Five unique haplotypes (Hap3-Hap7) were observed in the Uttarakhand population, whereas Hap2 and Hap8 were unique in Nepal and J&K populations, respectively (Fig. 2). The median-joining network of all haplotypes of six musk deer species strongly indicated the presence of geospatial population structures in *M. cupreus* and *M. leucogaster*, whereas weak structuring was observed in *M. berezovskii* and *M. chrysogaster*. The Bayesian phylogenetic result showed that the samples from Uttarakhand clustered with samples of KMD of J&K and submitted sequences from Nepal, and formed a separate clade (PP∼100%); while *M. moschiferus* formed the basal clade (Fig. 3). The BI tree topology indicated *M. cupreus* and *M. moschiferus* have evolved earlier than *M. chrysogaster, M. anhuiensis, M. leucogaster* and *M. berezovskii*.

**Figure 2.**
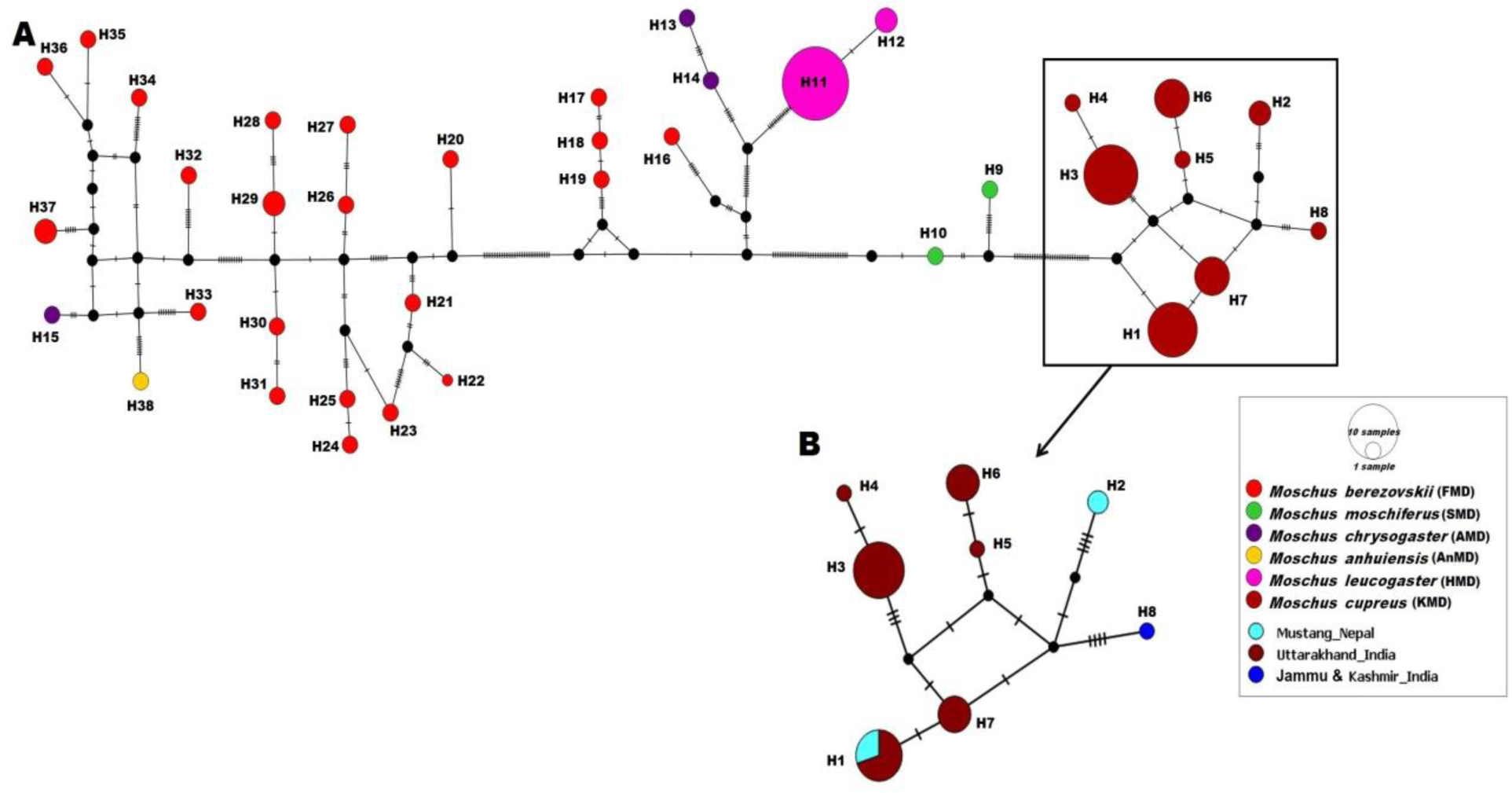
Median-joining (MJ) network based on the mtDNA control region of musk deer. The highlighted portion represents the sharing of haplotypes (H1-H8) of Kashmir musk deer (*M. cupreus*) from Nepal, Jammu and Kashmir and Uttarakhand, India.

**Figure 3.**
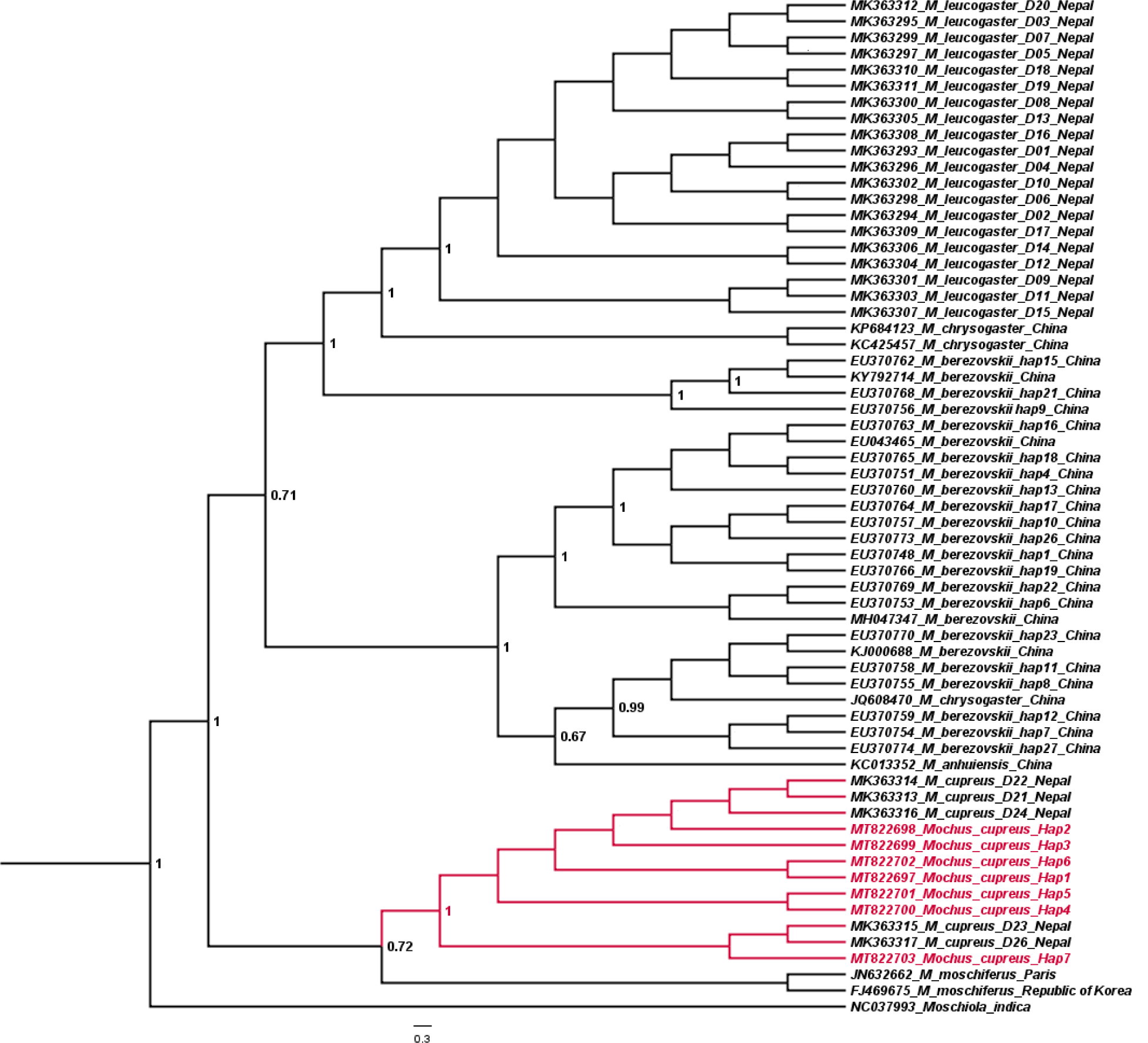
Bayesian (MCMC) consensus tree of musk deer based on the mtDNA control region. Posterior values are provided at their respective nodes. The *Moschiola indica* (NC037993) was used as the outgroup. The clade of Kashmir musk deer (*M. cupreus*) is highlighted in red color.

The mean pairwise genetic distance was calculated within and between the groups of musk deer species available in GenBank and with our generated data. The result indicated that the sequences of *M. cupreus* from Nepal were genetically similar to the J&K and Uttarakhand population with low sequence divergences estimates between the group (1%) and within species group (0.8%). Among the musk species, *M. cupreus* was found to be close to *M. moschiferus* (8.8-9.0%) followed by *M. leucogaster* (10%); whereas the maximum genetic difference of *M. cupreus* was observed with *M. berezovskii* (10.9%) (Table 1). High intraspecies sequences divergences were observed in *M. chrysogaster* (6.3%) and *M. berezovskii* (4.7%). The high sequence divergences and weak genetic clustering within the *M. chrysogaster* group raised concern on the authenticity of the complete mitogenome sequence (JQ608470) as *M. chrysogaster* by Yang et al., (2013) that clustered instead in the clade of *M. berezovskii* (Fig. 2 and 3), whereas the sequences of *M. berezovskii* also formed two different clusters. The high sequences divergence in *M. berezovskii* creates ground for comprehensive research to enable proper lineage confirmation.

**Table 1.** Genetic p-distance of the mtDNA CR of the genus *Moschus* are represented below the diagonal and standard error values are shown above the diagonal.

| Species | 1 | 2 | 3 | 4 | 5 | 6 | 7 |
| --- | --- | --- | --- | --- | --- | --- | --- |
| 1 *M. cupreus* (Nepal) |  | 0.003 | 0.012 | 0.013 | 0.013 | 0.014 | 0.012 |
| 2 *M. cupreus* (J&K;UK,India) | 0.010 |  | 0.012 | 0.013 | 0.013 | 0.014 | 0.013 |
| 3 *M. moschiferus* | 0.088 | 0.090 |  | 0.013 | 0.012 | 0.013 | 0.012 |
| 4 *M. leucogaster* | 0.100 | 0.101 | 0.094 |  | 0.009 | 0.013 | 0.011 |
| 5 *M. chrysogaster* | 0.103 | 0.102 | 0.095 | 0.047 |  | 0.012 | 0.008 |
| 6 *M. anhuiensis* | 0.107 | 0.112 | 0.100 | 0.090 | 0.063 |  | 0.007 |
| 7 *M. berezovskii* | 0.109 | 0.111 | 0.098 | 0.093 | 0.071 | 0.046 |  |

The presence of KMD was recently reported from Mustang, Nepal, which was previously believed to be restricted to the Himalayan region of Kashmir, Pakistan and Afghanistan (Grubb et al., 2005; Singh et al., 2019). Based on our genetic results, we confirm the distribution of KMD from J&K to the UK region of KWLS (1025–3662 m) and NDBR (1,800–7,817 m). The prediction based on climate refugia and habitat suitability mapping for KMD supported the probability of occurrence in the belt of high Himalaya region that stretches from central Nepal to the north-west of India including Uttarakhand and Himachal Pradesh, reaching Afghanistan through the Kashmir Region of India and Pakistan (Singh et al., 2020). In addition, as per the IUCN database, the Western Himalayan region of UK also holds the population of *M. leucogaster* and *M. chrysogaster*. Therefore, we suggest extensive sampling with ecological data and photographic evidence for the identification and confirmation of the distribution limits of musk deer species. All the species of musk deer should be considered as distinct Evolutionary Significant Units (ESUs), which require long term monitoring and special management attention. Proper knowledge about species distribution is important to build effective laws for their protection and conservation management.

## Conclusions

Therefore, in this study, we report the first distribution record of KMD (*M. cupreus*) from KWLS and NDBR in UK state of India using the hypervariable segment of the mtDNA CR. Our study provides the baseline evidence confirming the presence of KMD in Uttarakhand state of India which will be helpful in the reassessment of the species’ geographical distribution and also provide information to prepare effective conservation management strategies for the highly endangered species of musk deer. We recommend revision of the distribution range of KMD in the IUCN database to delineate the geographic boundaries for effective in-situ and ex-situ strategies for musk deer. A comprehensive ecological and molecular study is required with high throughput sequencing as well as microsatellite markers to understand the population dynamics of the musk deer species as well as molecular tracking of confiscated items in wildlife trade. A collaborative study from all range countries of musk deer species is vital for population and distribution assessment of this species.

## Acknowledgment

We thank Dr. Dhananjai Mohan, Director, Dr. Y.V. Jhala, Dean, and Dr. G.S. Rawat (former Dean and Director), WII for their support. We thank the State Forest Departments of Jammu and Kashmir and Uttarakhand for forwarding biological samples for the forensic examination to Wildlife Forensic and Conservation Genetics (WFCG) Cell, WII. We acknowledge the assistance of Mr. A. Madhanraj, WFCGC during this study.

---

**Supplementary Table ST 1.** Details of GenBank accession number used in phylogenetic analysis.

| **Species** | **Localities** | **Accession Number** | **Reference** |
| --- | --- | --- | --- |
| *M. moschiferus* | Republic of Korea | FJ469675 | Jang, et al., 2010 |
| *M. moschiferus* | Paris | JN632662 | Hassanin, et al.,2012 |
| *M. chrysogaster* | China | KC425457 | Wang et al., 2013 |
| *M. chrysogaster* | China | KP684123 | Pan et al.,2015 |
| *M. chrysogaster* | China | JQ608470 | Yang et al.,2013 |
| *M. anhuiensis* | China | KC013352 | Zhu et al.,2012 |
| *M. leucogaster* | Nepal | MK363293 | Singh et al., 2019 |
| *M. leucogaster* | Nepal | MK363294 | Singh et al., 2019 |
| *M. leucogaster* | Nepal | MK363295 | Singh et al., 2019 |
| *M. leucogaster* | Nepal | MK363296 | Singh et al., 2019 |
| *M. leucogaster* | Nepal | MK363297 | Singh et al., 2019 |
| *M. leucogaster* | Nepal | MK363298 | Singh et al., 2019 |
| *M. leucogaster* | Nepal | MK363299 | Singh et al., 2019 |
| *M. leucogaster* | Nepal | MK363300 | Singh et al., 2019 |
| *M. leucogaster* | Nepal | MK363301 | Singh et al., 2019 |
| *M. leucogaster* | Nepal | MK363302 | Singh et al., 2019 |
| *M. leucogaster* | Nepal | MK363303 | Singh et al., 2019 |
| *M. leucogaster* | Nepal | MK363304 | Singh et al., 2019 |
| *M. leucogaster* | Nepal | MK363305 | Singh et al., 2019 |
| *M. leucogaster* | Nepal | MK363306 | Singh et al., 2019 |
| *M. leucogaster* | Nepal | MK363307 | Singh et al., 2019 |
| *M. leucogaster* | Nepal | MK363308 | Singh et al., 2019 |
| *M. leucogaster* | Nepal | MK363309 | Singh et al., 2019 |
| *M. leucogaster* | Nepal | MK363310 | Singh et al., 2019 |
| *M. leucogaster* | Nepal | MK363311 | Singh et al., 2019 |
| *M. leucogaster* | Nepal | MK363312 | Singh et al., 2019 |
| *M. berezovskii* | China | EU370748 | Peng et al.,2009 |
| *M. berezovskii* | China | EU370751 | Peng et al.,2009 |
| *M. berezovskii* | China | EU370753 | Peng et al.,2009 |
| *M. berezovskii* | China | EU370754 | Peng et al.,2009 |
| *M. berezovskii* | China | EU370755 | Peng et al.,2009 |
| *M. berezovskii* | China | EU370756 | Peng et al.,2009 |
| *M. berezovskii* | China | EU370757 | Peng et al.,2009 |
| *M. berezovskii* | China | EU370758 | Peng et al.,2009 |
| *M. berezovskii* | China | EU370759 | Peng et al.,2009 |
| *M. berezovskii* | China | EU370760 | Peng et al.,2009 |
| *M. berezovskii* | China | EU370762 | Peng et al.,2009 |
| *M. berezovskii* | China | EU370763 | Peng et al.,2009 |
| *M. berezovskii* | China | EU370764 | Peng et al.,2009 |
| *M. berezovskii* | China | EU370765 | Peng et al.,2009 |
| *M. berezovskii* | China | EU370768 | Peng et al.,2009 |
| *M. berezovskii* | China | EU370769 | Peng et al.,2009 |
| *M. berezovskii* | China | EU370770 | Peng et al.,2009 |
| *M. berezovskii* | China | EU370773 | Peng et al.,2009 |
| *M. berezovskii* | China | EU370774 | Peng et al.,2009 |
| *M. berezovskii* | China | KJ000681 | Yang et al.,2014 |
| *M. berezovskii* | China | KJ000688 | Yang et al.,2014 |
| *M. berezovskii* | China | KY792714 | Su et al.,2018 |
| *M. berezovskii* | China | MH047347 | Yang et al.,2018 |
| *M. berezovskii* | China | EU043465 | Peng et al.,2009 |
| *M. cupreus* | Nepal | MK363313 | Singh et al, 2019 |
| *M. cupreus* | Nepal | MK363314 | Singh et al, 2019 |
| *M. cupreus* | Nepal | MK363315 | Singh et al, 2019 |
| *M. cupreus* | Nepal | MK363316 | Singh et al, 2019 |
| *M. cupreus* | Nepal | MK363317 | Singh et al, 2019 |

